# Circularly shifted filters enable data efficient sequence motif inference with neural networks

**DOI:** 10.1101/312959

**Authors:** Christopher F. Blum, Markus Kollmann

**Affiliations:** Institute for Mathematical Modeling of Biological Systems, Heinrich-Heine University of Düsseldorf, 40225 Düsseldorf, Germany

## Abstract

**Motivation:** Nucleic acids and proteins often have localized sequence motifs that enable highly specific interactions. Due to the biological relevance of sequence motifs, numerous inference methods have been developed. Recently, convolutional neural networks (CNNs) achieved state of the art performance because they can approximate complex motif distributions. These methods were able to learn transcription factor binding sites from ChIP-seq data and to make accurate predictions. However, CNNs learn filters that are difficult to interpret, and networks trained on small data sets often do not generalize optimally to new sequences.

**Results:** Here we present circular filters, a novel convolutional architecture, that contains all circularly shifted variants of the same filter. We motivate circular filters by the observation that CNNs frequently learn filters that correspond to shifted and truncated variants of the true motif. Circular filters enable learning of non-truncated motifs and allow easy interpretation of the learned filters. We show that circular filters improve motif inference performance over a wide range of hyperparameters. Furthermore, we show that CNNs with circular filters perform better at inferring transcription factor binding motifs from ChIP-seq data than conventional CNNs.

## 1 Introduction

A fundamental property of biological macromolecules such as DNA, RNA and proteins is their ability of highly specific interactions. These high specificities are often associated with localized motifs in the primary structure of these macromolecules. Intricate motif variations, indirect effects on binding specificity and noisy data make motif inference from high-throughput data a challenging task [1, 2, 3, 4, 5].

As DNA, RNA or proteins participate in almost all processes that have biotechnological or biomedical relevance, numerous methods have been developed to infer motifs and binding specificities from high-throughput data sources [5]. These methods range from derivatives of nucleotide frequency counting procedures such as *k*-mer and position weight matrix (PWM) methods to, most recently, convolutional neural networks (CNNs) [6, 7]. Unlike PWM methods, CNNs do not require aligned input sequences because they convolve input sequences with multiple sliding weight matrices called filters. Although this sliding operation resembles *k*-mer methods, CNNs do not rely on predefined *k*-mers. Instead, they learn the weight matrices by maximizing an objective function. This function is either a distance measure between observed and predicted fluorescence intensities or read counts or class-conditional probabilities of labelled sequences. While position weight matrices allow intuitive interpretation of the weight matrices in terms of information content with respect to a background model, CNNs distribute a motif’s distributional information across the multiple filters in a non-trivial way, preventing immediate interpretation.

Convolutional neural network architectures have achieved state of the art performance at predicting fluorescence intensities derived from protein binding microarray (PBM) data [6]. It has been reported, however, that training these models with gradient descent methods can be sensitive to weight initialization, impairing generalization ability [7]. As Stochastic Gradient Langevin Dynamics (SGLD) has been proposed as a general-purpose method for approximate bayesian inference that can strongly reduce the risk of overfitting, we utilize SGLD for motif inference with CNNs [8].

Deep neural networks typically require large amounts of training data. Due to the noise of PBMs and ChIP-seq derived data, the number of positive training examples is often limited. A CNN-based motif inference method, DeepBind, addresses this problem by artificially increasing the number of negative training examples with data augmentation, but still relies on a sufficient amount of positive training examples, that is, sequences with high fluorescence intensities or read counts. [6, 9].

Here, we present a neural network algorithm that utilizes a novel convolutional architecture called circular filters that enables efficient data utilization and easy interpretation of the learned filters. First, it is shown that conventional CNN filters often contain shifted and truncated motifs. We then introduce circular filters as a natural solution to this problem and show that motif inference improves for a wide range of hyperparameters, allowing better inference when data is scarce. Finally, we demonstrate that a CNN with circular filters and SGLD yields accurate predictions for ChIP-seq derived data, resulting in the state of the art algorithm for motif inference from smaller-sized data sets.

## 2 Results

### 2.1 Convolutional neural networks frequently learn truncated motifs

While studying motif inference with CNNs, we observed that the learned filter weights frequently did not correspond to the complete motif. Instead, the filters often contained truncated and shifted versions of the motif (Fig. 1a). To quantify this behaviour, we conducted simulations in which a known motif had to be inferred from a set of short sequences. Then, it was counted how frequently the filter contained to a shifted version of the motif. We found that CNNs learned shifted and truncated motifs more frequently than the true motifs (Fig. 1b).

**Figure 1:**
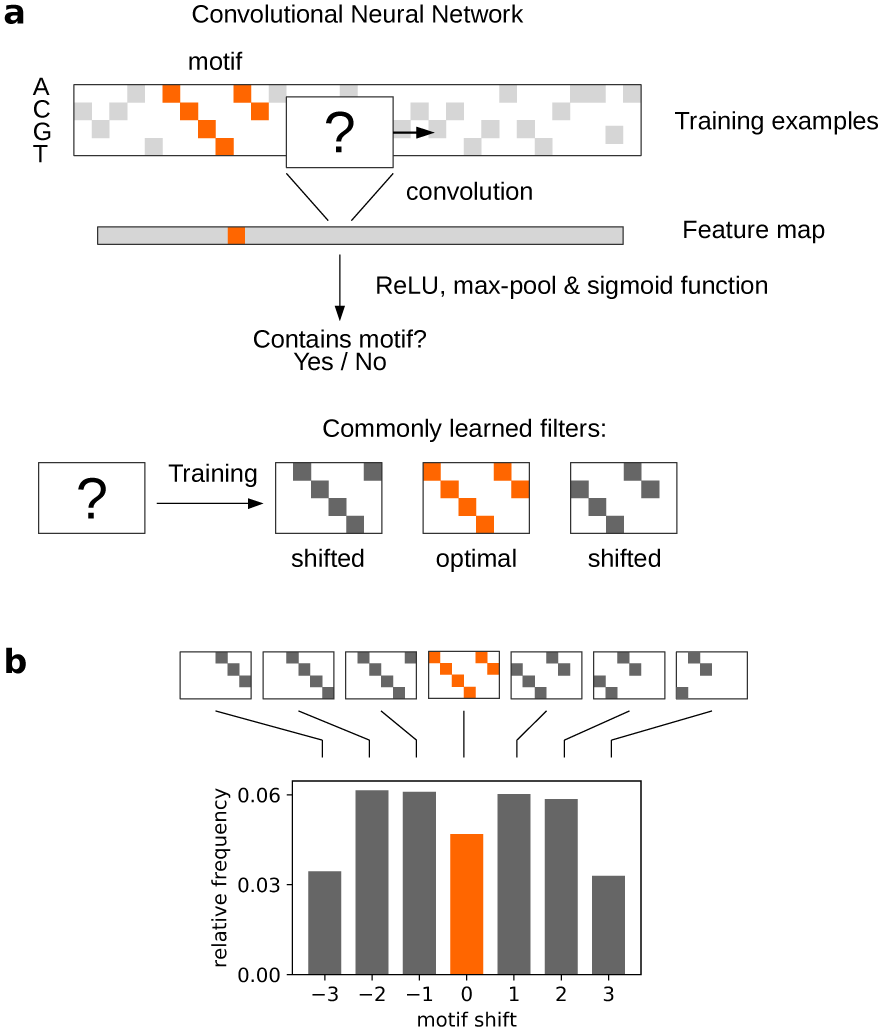
**a)** Illustration of a conventional convolutional neural network (CNN). The network is trained to discriminate between sequences with and without motifs in a supervised manner. Comparison of the learned filters with the actual motif reveals that they often contain a truncated and shifted version of the motif. **b)** Shifted motif variants are being learned more frequently than the correct, unshifted motif by conventional CNNs. Bars show relative frequency of shifts derived from CNN filters that were trained on 4096 different data sets corresponding to all possible 6-mers, with 100 positive examples in each data set. The motifs pointing to the bars are for illustration purposes only and do not represent the actually learned motifs.

### 2.2 Circular filters improve robustness of sequence motif inference for simualted data

This observation motivated us to develop a novel convolutional kernel that already contains all circularly shifted variants of the same underlying filter, which we refer to as circular filters (Fig. 2a). If one of the filters in this kernel learns a shifted motif, there is another filter variant that is able to learn the full motif, provided the filters have at least the size of the motif. Based on the feature map with the largest activation, the filter variant that contains the non-shifted motif can be recovered. In simulations we found that CNNs with circular filters rarely learn shifted motifs (Fig. 2b). When trained on 100 positive examples to infer motifs of 6 nucleotides length that are randomly located in a 40 nucleotide long sequence, a CNN with circular filters learned the correct motifs 5.5 times more often than a CNN without circular filters (Table 1). Even when the CNN with circular filters was trained with only 5 positive examples, the correct motif was found 1.9 times more often (*p* < 10^−27^).

**Figure 2:**
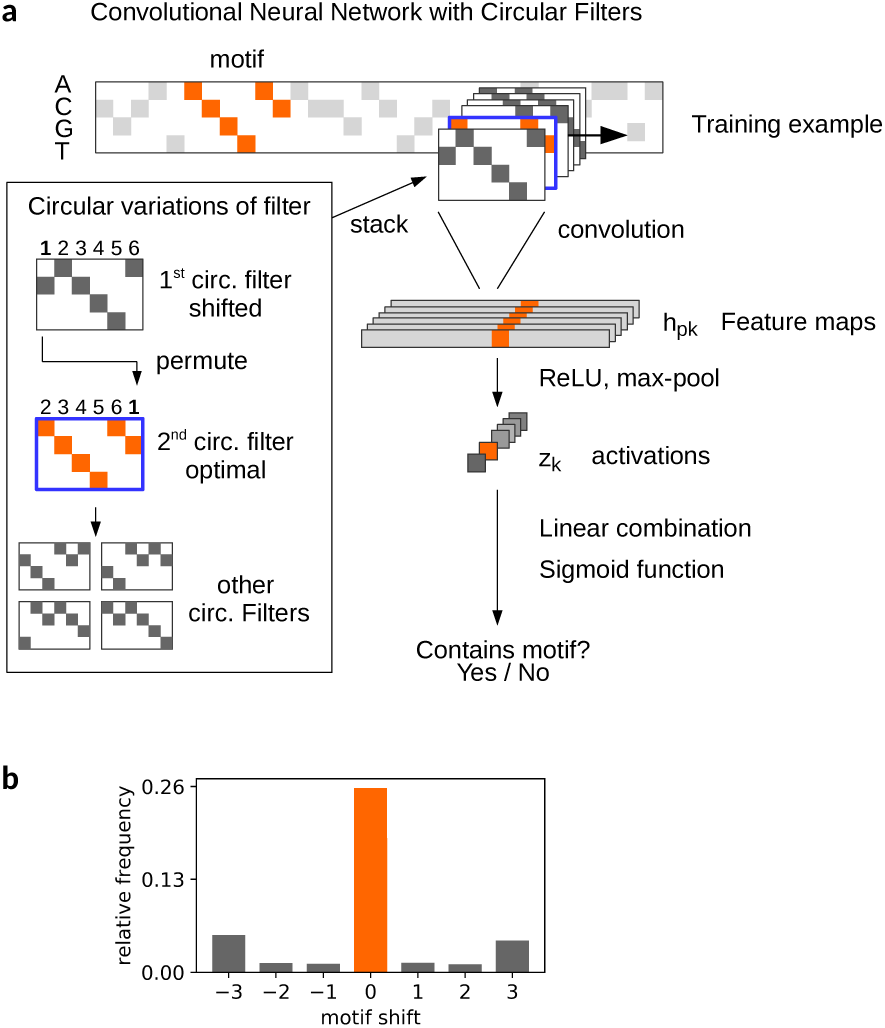
**a)** Illustration of a convolutional neural network with circular filters. The circular filter consists of all possible circular variants of the same underlying filter. Convolution with one circular filter of length *N* yields *N* feature maps that are linearly combined after max-pooling. **b)** The correct motif is learned more frequently than shifted motifs. Bars show relative frequency of shifts derived from CNN filters that were trained on 4096 different data sets corresponding to all possible 6-mers, with 100 positive examples in each data set.

We quantified the effect of circular filters on motif recognition by comparing network architectures with and without circular filters for a variety of hyperparameter combinations. These included the number of positive training examples, L2-regularization strength and the amount of noise injected into parameter updates. As the original architecture with circular filters (Fig. 2a) forms a linear combination of the activations which increases the number of parameters compared to the architecture without circular filters (Fig. 1a), we also designed two architectures without additional parameters. These architectures use a simple sum or max-out of the activations [10]. Overall, both the CNN with circular filters and the CNN with circular filters and sum of activations performed significantly better than the regular CNN in 74% and 61% of all cases, respectively (*p* ≤ 10^−5^). The CNN with circular filters and max-out however performed significantly worse than the regular CNN in 56% of cases (*p* ≤ 10^−5^). SGLD did not significantly improve performance, but too much noise injection into parameter updates significantly reduced performance (Table 2). Adjustment of L2-regularization strength significantly improved motif inference for all methods (Table 3).

### 2.3 Circular filters lead to state of the art performance for motif inference

To investigate if circular filters improve inference of biological motifs, we trained a CNN with circular filters to infer transcription factor binding sites from ChIP-seq data [11]. The area under the receiver operator characteristic (AUROC) on evaluation data sets was used as a performance measure. The state of the art method for motif inference, DeepBind, was used as a reference [6]. Performance of both methods was assessed for different filter sizes, numbers of filters, and number of positive training examples.

We found that the CNN with circular filters led to AUROC values greater than those of DeepBind for a variety of parameter combinations (Fig. 3, Table 4). The difference in performance was especially large when only a single filter was used and 500 positive training examples were available (Fig. 4). With more and larger filters and less positive training examples, the difference in performance was generally smaller.

**Figure 3:**
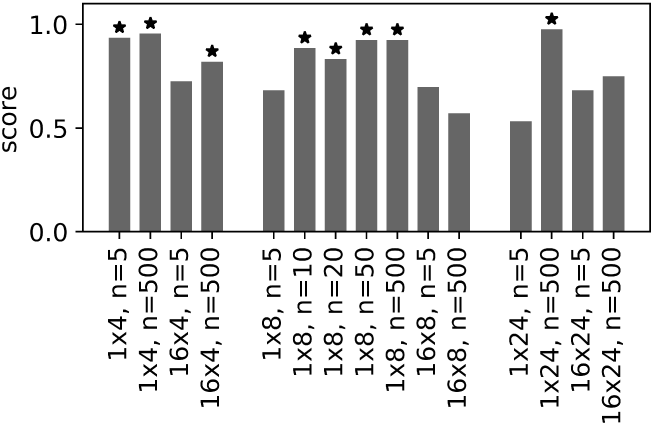
Performance comparison between a CNN with circular filters and DeepBind on ChIP-seq data published by the ENCODE consortium. The score indicates the ratio of times that the CNN with circular filters led to AUROC values larger than those obtained the state of the art method for inferring sequence motifs. Labels indicate number and size of filters as well as the number of positive examples, e.g. 1 × 4, *n* = 5 means that one filter of length 4 was used on a data set with 5 positive examples. Asterisks indicate p-values below 10^−5^ (Binomial test).

**Figure 4:**
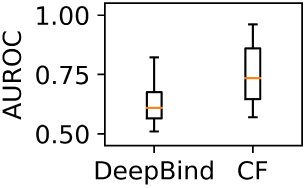
Circular filters help to infer and predict transcription factor binding sites. Comparison based on ChIP-seq data sets published by the ENCODE consortium. Both algorithms were trained on 500 examples and used 1 filter of length 8 nucleotides. Box-plot whiskers show 2.5% and 97.5% quantiles.

Furthermore, we found that the CNN with circular filters produced AUROC values above 0.5 more often than DeepBind (Table 4). Especially when only one filter was used, DeepBind yielded AUROC values smaller than 0.5 both when either 5 or 500 positive training examples were provided. In contrast, the CNN with circular filters displayed this behaviour only when 5 positive training examples were available.

## 3 Methods

### 3.1 Investigated network architectures

### 3.2 Comparing learned filters and true motifs

Synthetic data sets were generated for all 4096 6-mers. Each data set consisted of either 5 or 100 positive and 10, 000 negative examples. The positive examples were created by embedding a 6-mer in random sequences of 40 nucleotides length each. The set of 10, 000 negative examples was created by randomly sampling (with replacement) from the set of positive examples and then randomly shuffling the nucleotide order of each sequence. All sequences were converted to a one-hot representation to be used as input by the network.

Two network architectures were investigated: the regular CNN and the CNN with circular filters (Fig. 1a and Fig. 2a, respectively). The networks were trained for 10, 000 steps at a learning rate of 0.01.

The learned filters were converted to a one-hot representation indicating the largest weight at each filter position and then and compared to the original 6-mer motif. In this comparison, it was checked if the learned filters corresponded to shifted and truncated versions of the original motif. Specifically,

- shifts −3, −2 and −1: it was checked if the *l* rightmost nucleotides of the filter were equal to the *l* leftmost nucleotides of the motif hidden in the sequences, with *l* ∈ 3, 4, 5,
- shift 0: it was checked if all 6 nucleotides of the filter were equal to the complete motif hidden in the sequences
- shifts 1, 2, 3: it was checked if the *l* leftmost nucleotides of the filter were equal to the *l* rightmost nucleotides of the motif hidden in the sequences, with *l* ∈ 3, 4, 5.

The number of times, *k*, with which shifted motifs were learned from the *n* = 4096 data sets was tested for statistical significance using the Binomial distribution *B_n,p_*(*k*), where 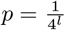 is the probability to observe a motif of length *l* by chance. Since some of the resulting p-values were too small to be calculated, the smallest *k* necessary for a p-value less than or equal to 10^−5^, *k*_signif_, was calculated instead. For example, the value of the leftmost bar in the barplot in Fig. 1b at 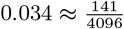 is significant because a shift of −3 was observed 141 times and *k*_signif_ was 100.

To assess whether the CNN with circular filters learned unshifted motifs significantly more often than the regular CNN, we modelled the null hypothesis with a binomial distribution *B_n,p̂_*(*k*), with *n* = 4096 and *p̂* the estimated probability of the regular CNN learning an unshifted filter, and *k* the number of times the CNN with circular filters learned an unshifted motif.

### 3.3 Influence of circular filters on motif recognition

To investigate the effect of circular filters and hyperparameters on motif inference, we trained and evaluated all network architectures on simulated data for a variety (grid) of hyperparameter combinations and then used statistical tests to identify significant effects.

We generated synthetic data sets of labelled sequences with or without a motif for training and evaluation. First, 32 random motifs of 6 nucleotides length were generated. For the training data, three data sets corresponding to the number of positive training sequences (5, 50 and 500) were generated for each motif. Depending on this number, the corresponding number of random nucleotide sequences of 40 nucleotides length was generated, and a motif was then placed at a random position in each of these sequences. These sequences corresponded to the positive examples. Negative examples were generated by randomly sampling 10, 000 times from the positive examples and then randomly shuffling the nucleotide order. Evaluation data sets were created for each of the 32 motifs and consisted of 1000 positive and 1000 negative examples. All sequences were converted to one-hot coding.

Networks were trained for 40000 training steps at a learning rate of 0.01. Mini-batches consisted of randomly (with replacement) selected 10 positive examples and 10 negative examples. The cross-entropy between sequence labels and predicted sequence labels was used as objective function and was minimized using SGLD [8]. AUROC values were calculated based on predictions on the evaluation data sets.

To identify significant effects of circular filters, SGLD and L_2_-regularization, the number of times that a specific architecture or hyperparameter value overall resulted in higher AUROC values than a reference was calculated. To test these numbers for significance, we approximated cumulative distribution functions (CDF) by bootstrapping. The reason is that the AUROC values could not be assumed to be independently distributed: the same 32 motifs were used to investigate all hyperparameter combinations, and some motifs might have been generally easier to infer than others. Hence we had to assume the existence of correlations between the performances of different hyperparameter combinations applied to the same motif. To destroy such correlations, we created bootstrap samples from the obtained AUROC values by randomly drawing AUROC values corresponding to different motifs. To approximate the cumulative distribution functions with sufficient accuracy, we used 10^5^ bootstrap samples. The smallest and largest approximated CDF values that could be resolved with this method were 10^−5^ and 1 − 10^−5^.

Architectures and hyperparameters were compared as follows and for the following values: The architectures with circular filters were compared to the architecture without circular filters, the effect of SGLD noise variance scaling factor was investigated relative to a scaling factor of zero (Table 2), and the effect of L_2_-regularization strength was investigated relative to a strength of zero (Table 3). Values of 0.1, 1.0, 10 and 100 were used for both the L_2_-regularization strength and SGLD noise variance scaling factor.

### 3.4 Performance comparison between DeepBind and SGLD using ENCODE data

The performance of DeepBind and the CNN with circular filters was evaluated for several combinations of filter length, number of filters and number of positive training examples (Fig. 3). Both algorithms were provided with exactly the same training and test data.

Data preprocessing is described elsewhere in more detail [6]. In short, input data was derived from 48 randomly selected transcription factor ChIP-seq experiments for which the data was published by the Encyclopedia of DNA Elements (ENCODE) Consortium [11]. These ChIP-seq data sets comprised of mapped sequence reads and read numbers for the most significant peaks. Regions around read peaks (starting 50 nucleotides before and ending 50 nucleotides after each peak) were extracted and sorted according to the read number in descending order. As the first 1000 sequences had the largest read numbers, it was assumed that they contained a binding site for the respective transcription factor. These sequences constituted the positive examples. For each ChIP-seq experiment, the first 500 even-numbered positive examples were allocated to the corresponding test data set. Depending on the setting, either the first 5, 10, 20, 50 or 500 odd-numbered sequences were allocated to the corresponding training data set.

Negative examples for the training and test data sets were created based on the corresponding positive examples. For the training data, 10, 000 sequences were randomly sampled with replacement from the positive examples. Then, the nucleotide order within the individual sequences was shuffled using dinucleotide shuffling, a method that shuffles the nucleotide order while preserving the frequency of neighbored nucleotides [12]. For the test data, two data sets were created by first copying all 500 positive examples and then shuffling the nucleotide using either random shuffling and dinucleotide shuffling.

Because our computational power was limited, we restricted our comparison to randomly selected 48 out of the 506 ChIP-seq data sets that were used in the comparison by the DeepBind authors.

Both DeepBind and the CNN with circular filters assigned a score to each sequence, indicating the likelihood that the sequence contained a motif. By assigning a score to each of the 500 positive and 500 negative examples in the test data sets, AUROC values could be calculated.

Before any further testing, only runs in which both CNN with circular filters and DeepBind had AUROC values greater than 0.01 were used. This was found to be necessary because for some runs, DeepBind assigned a class conditional probability of 0.5 to all positive and negative examples, resulting in AUROC values equal to 0.0. This behaviour has been reported before [7]. For each parameter combination and ChIP-seq experiment, the number and fraction how often CNN with circular filters had AUROC values above the corresponding DeepBind AUROC values was obtained and tested for significance (Fig. 3 and Table 4, respectively). The null hypothesis was modelled with a binomial distribution *B_n,p_*(*k*) with *n* the number experiments and *p* = 0.5. Because it can happen that neural networks do not converge to the global minimum during training, ultimately yielding AUROC values either close to or even below 0.5, a second statistic was calculated as well that disregarded such runs. For this statistic, only the runs in which both CNN with circular filters and DeepBind led to AUROC values greater than 0.5 were used to calculate the number of times CNN with circular filters yielded AUROC values greater than those of DeepBind.

#### DeepBind settings

The default DeepBind training parameters were used, except for an override of the filter length and the number of filters. Furthermore, the original routine for generating negative examples de novo was replaced by a routine to load the negative examples from the hard drive to ensure better comparability. To obtain a platform-independent instance of DeepBind, DeepBind was run in a Docker [13] container using Nvidia Docker. This was necessary because former attempts towards a platform-independent implementation [7] were still depending on a certain CUDA-version [14].

#### Settings for CNN with circular filters

Models were trained using a learning rate of 0.01 for 40, 000 training steps. We used a batch size of 20 consisting of 10 positive and 10 negative examples, all randomly sampled (with replacement) from the respective training data.

For hyperparameter tuning, the model was trained for 5 different regularization strengths. The training data was split up into a training and a validation data set (80% and 20% of the data, respectively). Every 50 steps, it was determined whether the error on the validation set was lower than before, in which case the current model was stored for later restoring. Then, the model corresponding to the regularization strength that yielded the lowest error on the validation set was used for evaluation on the test data set, for which the the model that yielded the lowest validation error was restored.

## 4 Algorithm

### 4.1 One-hot coding

A sequence Σ = Σ_1_,…,Σ_*L_S_*_ with elements Σ_*j*_ coming from an ordered set with limited finite cardinality Σ_*i*_ ∈ *D*, |*D*| = *N* can be represented as a *N* × *L_S_* one-hot coding matrix S with elements *S_ij_* that are equal to 1 if Σ_*j*_ is the i-th element in *D* and 0 otherwise (Fig. 5).

**Figure 5:**
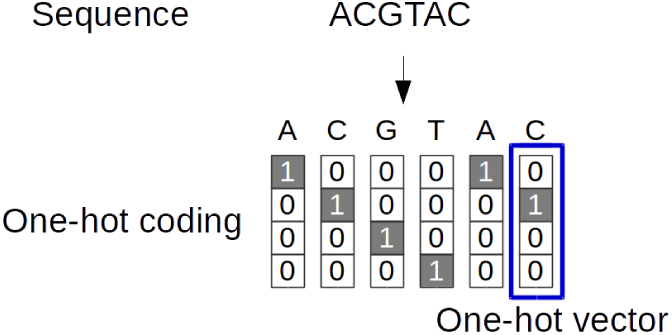
Illustration: a nucleic acid sequence represented as one-hot coding.

### 4.2 Formal definition of 1-dimensional convolution with one circular filter

The unpadded convolution of a one-hot coded sequence *S* with length *L_S_* and *N* features and one circular filter of length *L_F_* and *N* features to a hidden node *h_pk_*, *p* ∈ {1,…, *L_S_* − *L_F_* + 1}, *k* ∈ {1,…, *L_F_* }, can formally be defined as:

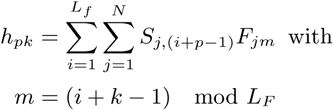

### 4.3 Formal definitions of the network architectures with circular filters

We model the class-conditional probability *p*(*C*_motif_*|S_n_*) that a sequence with one-hot coding *S_n_* belongs to class *C*_motif_ (“contains motif”) as a convolutional network. Input to the network is a *N* × *L_S_* one-hot coding *S_n_*, where *L_S_* is the length of the sequence and *N* is the number of features (that is, the number of different nucleotides or amino acids). The input sequence *S_n_* is convolved without padding, followed by a Rectifying linear unit (ReLU) and max-pooling to yield activations *z* that are mapped onto the class-conditional probabilities depending on the architecture. The objective function was the batch-averaged cross-entropy between predicted and observed classes. In the following, the *N* × *L_F_* matrix *F* is the convolutional filter of length *L_F_*, *w_k_* or *w* are weights and *b* is a bias, and 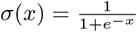 denotes the sigmoid function.

The CNN without circular filters (Fig. 1a) is a chain of functions in the following order:

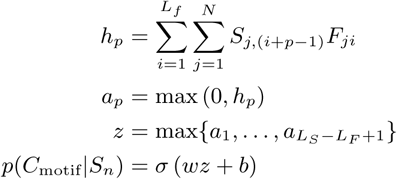

The CNN architectures with circular filters used here all convolve the input with circular filters but differ in the operation applied to the activations *z_k_*, specifically:

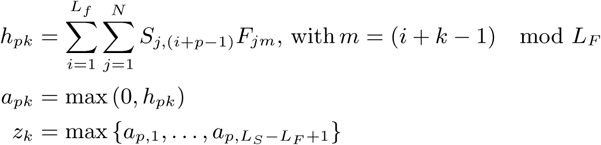

For the convolutional neural network with circular filters (Fig. 2a), we then have:

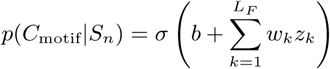

Whereas for the convolutional neural network with circular filters and sum of the activations, we have:

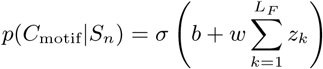

Finally, for the convolutional neural network with circular filters and max-out, we have:

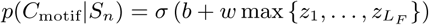

## 5 Discussion

### 5.1 Motif inference differs from deep learning disciplines

We showed in a simple simulation that a shallow convolutional neural network architecture frequently learns non-optimal filters that correspond to shifted and truncated versions of the underlying inferable pattern. The networks were trained with stochastic gradient descent, indicating that the non-optimal filters correspond to locally optimal variable combinations. This is in agreement with the well-studied behavior of gradient descent optimization which is highly sensitive to initial conditions and prone to local optima. The local optima observed here seem to be more difficult to escape compared to other machine learning disciplines where CNNs are applied.

### 5.2 Circular filters enable robust motif inference

The filter weights of conventional CNNs are trained by starting from random seeds. For motif inference, this seed determines at which position in the filter the inferred motif will be developed during training. This means that it can happen that only one side of a motif can be learned because an edge of the filter has been reached. With circular filters, there are no edges and the position from where the motif develops does not matter anymore.

This leads to the following consideration on a motif inference problem in which one filter shall be learned whose length is equal to the motif. For a conventional CNN, there can be only one optimal filter, corresponding to one global optimum in parameter space. For a CNN with circular filters, however, there can be at most *N* global optima (with *N* denoting filter length). Our results suggest a linear relationship between filter length and number of global optima (Fig. 3). This should affect the probability of gradient descent optimization converging to a global optimimum and may explain why the CNN with circular filters performed better than the regular CNN despite near identical number of parameters.

It has been observed before that that slight modifications in network structure alongside negligibly more parameters can substantially improve inference performance [15, 16]. These findings have demonstrated the importance of hard-wired prior knowledge about the underlying problem. Also circular filters can be regarded as hard-wired prior knowledge: sequence motifs are locally correlated data features that require convolution to be learned and optimization with gradient descent requires sufficient space to develop the motif from initial weights.

Since the three investigated architectures with circular filters only differed in the function applied to the activations, this must be the reason for the observed performance differences. For the CNN with circular filters and max-out, backpropagation of the classification error can only occur to the circular filter variant that led to the maximum activation. After applying the parameter updates, it can happend that another circular filter variant leads to the maximum acitivation in the next training step.

This may complicate parameter optimization, explaining the lower performance compared to the architecture without circular filters. For the other architectures, the classification error can backpropagate to all filter variants, allowing the motif to be learned in any of the circular filters. However, also filter variants that do not contain the inferable pattern can contribute to the classification error, which can be harmful if some training sequences also randomly contain other patterns. For the CNN with circular filters, it can be adjusted by a linear combination how much the filter variants contribute to the classification, which is not the case for the CNN with circular filters and sum of activations, which may explain the difference in performance.

### 5.3 Deep models are not necessary for modeling TF-DNA specificities

It has been implied that deep neural networks were necessary to model transcription factor specifitities because a convolutional neural network architecture achieved state of the art performance at predicting probe intensities derived from protein binding microarray data [6]. Although neural networks are complex function approximators and a deep CNN with 152 layers achieved state of the art performance at classifying images, it was demonstrated that deeper architectures actually perform worse at learning sequence motifs than architectures with one layer [17, 16, 7].

A likely reason is that biological sequences are not composed of complex hierarchies of patterns such as those in images. In fact, mutually exclusive sequence patterns or spatial relationships can already be modeled with two layers, and to the best of our knowledge, no transcription factor has yet been found which binds to mutually exclusive motifs. This likely makes truly deep architectures unnecessary and possibly deleterious because more parameters need to be trained. Also, most transcription factors bind to motifs of 30 nucleotides length, a size that can be captured with simple convolutional filters [18]. Because of the aforementioned reasons, it is unsurprising that the discriminative implementation of DeepBind has only two layers.

### 5.4 Summary

When applied to biological data, a CNN with circular filters performed at least as good as the current state of the art algorithm for several combinations of filter number and size. Even for small sample sizes, the CNN with circular filters was able to infer correct motifs more often than a CNN without circular filters trained with 20 times more examples. Overall, our findings show that circular filters enable more efficient use of data for sequence motif inference.

